# HemeFinder: a Computational Predictor for Heme-Binding Sites in Proteins

**DOI:** 10.1101/2025.04.14.648773

**Authors:** Laura Tiessler-Sala, Raúl-Fernández, Jean-Didier Maréchal

## Abstract

HemeFinder has been developed to predict heme-binding sites in natural and heme-dependent de novo enzymes. This tool relies on the structural and physicochemical characteristics of heme-binding sites, including shape, residue composition, and geometric descriptors. HemeFinder benchmarks more than 94% accuracy in identifying the correct heme location, considering the complete set of solutions, and 72% accuracy for the proper location with the correct iron-coordinating residues among the three best-ranked solutions. HemeFinder performs within seconds for monomeric systems and takes minutes for larger multimeric ones, demonstrating that its speed does not compromise its performance. An illustrative case of its potential is provided. HemeFinder is applied to the heme carrier protein 1 (HCP1), a transmembrane protein involved in heme recruitment in evolved organisms, for which no ligand-bound structures have been revealed. HemeFinder provides a relevant prediction of the binding of porphyrin and, when combined with protein-ligand docking, offers the first evidence of low-energy Heme-HCP1 complexes and unveils possible heme pathways. HemeFinder is an interesting, fast, and accurate tool for identifying heme-binding sites in proteins.

Source code, documentation, and data are available at https://github.com/laura-tiessler/hemefinder and ESI.

## INTRODUCTION

Understanding the nature of the interactions between inorganic elements and their biological partners is crucial for understanding the origins of life and exploring new biotechnological routes. The intersection between biology and inorganic chemistry presents challenges in assessing the influence of each partner in the recognition process. Among the most complex questions that could be addressed is the study of metal-mediated binding processes, particularly those involving essential metallic moieties like heme.

Heme is one of the most ubiquitous metal containing ligands in nature, and with a high biological relevance in several well-studied biological functions, including storage and transport of O_2_ (myoglobin and hemoglobin), electron transfers (cytochromes) and redox catalysis in enzymes (cytochromes P450, peroxygenases or oxidases)^1,2^. Additionally, heme contributes to complex regulation functions, including gas sensing and regulation of transcription.^3,4^ Despite its importance, the binding of heme to proteins and its prediction has not been extensively studied. The ability to predict and identify heme-binding sites is essential for understanding better natural heme-binding processes, but it can also provide insight in the design of heme-based artificial metalloenzymes (ArM).

Around 2010, several groups started to investigate heme-protein interactions in depth. Some studies focused on identifying the main structural motifs and interactions between heme and proteins, while others reported on how specific structural characteristics of heme-binding sites lead to different heme functions.^5,6^ Li *et al* studied the heme-binding site environment, pointing at the fact that heme-binding pockets contain mainly aromatic and non-polar residues. This suggests that coordination of the iron by an electron-rich amino-acid side chain like cysteinate or histidine and that polar interaction of the proponiate moieties to with positively charged amino acids (mainly arginine and lysine) are not the most predominant aspects in binding (though still being important for the catalytic function and the correct orientation of the prosthetic group). Moreover, a visual comparison of apo-holo heme-binding structures revealed that in most cases proteins suffer very small conformational changes upon heme binding; a phenomenon particularly true for transient heme-binding processes like those required in heme transport.^7^ Based on these works and as experimental determination is time-consuming and expensive, development of computational methods for predicting heme-binding sites have emerged over the past years.

Two main approaches exist for predicting heme-binding sites. The first involves predictors based purely on sequence information, as in the case of SCMHB^8^ or Xiong et al predictor.^9^ The former only relies on sequence information, while the latter combines sequence, evolutionary and physicochemical properties. More recently, the SeqD-HBM algorithm was developed based on experimental analysis of heme-peptides, which lead to a set of sequence features for predicting heme motifs.^10^ This algorithm is now implemented on HeMoQuest, a tool aimed to predict transient heme-binding sites.^11^

The second approach combines structural and sequence information. HemeBIND was the first to predict heme-binding residues based on supervised machine learning and integrating structural attributes like solvent accessibility or depth with sequence information like residue evolutionary conservation.^12^ The same group latter produced HemeNET, which also integrates residue interaction network.^13^ Another software, HEMEsPred also combines structural and sequence information to predict specific heme-binding sites, using an adaptive ensemble learning method to enhance the prediction.^14^ Recently, the ProFunc tool has been used to identify heme regulatory proteins.^15^

So far, heme predictors are sequence based or hybrid structure/sequence methods. Although these approaches are particularly relevant for natural enzymes, they can be a limitation for de novo systems which have not yet evolved for heme-binding processes. That would be the case of heme-based artificial metalloenzymes (ArM).

The aim of the current work is to provide a new software for detecting heme-binding sites based exclusively on the protein structure and the geometrical predisposition of heme-binding sites. Furthermore, due to the principles of this software, a novel functionality is implemented to contribute to the design of heme-based ArMs by predicting possible mutations that would increase heme affinity.

## COMPUTATIONAL METHODOLOGY

### Concept of the software

This program was conceptualized from the idea of BioMetAll^16^, which is a predictor of metal binding sites based only on the preorganization of the backbone. It uses three geometrical criteria from the backbone to make the predictions: the distance from the metal to the α-carbon (*dist(α-M))*, the distance between the metal and the β-carbon (*dist(β-M))* and the angle between the metal, the α-carbon, and the β-carbon (angle*(α-β-M))*. Despite the success of BioMetAll in predicting metal binding sites, it is not entirely suitable for detecting heme-binding sites because it does not consider the real dimension of the prosthetic group including the volume of the binding site, or the properties of the surrounding residues. Moreover, BioMetAll does not account for the possibility of axial coordination by either one or two amino acids.

Here, we present HemeFinder that maintains the central philosophy of BioMetAll, but includes additional descriptors of heme binding (**Figure 4.1**). HemeFinder aims to predict heme-binding sites considering only structural information in a fast approach relying on three assumptions: **1)** All heme-binding sites share common geometrical properties (volume and shape) for a well pre-organized binding site. **2)** The environment of the heme-binding site has certain residue composition and physicochemical properties. **3)** For those amino acids that coordinate the iron at the distal sites, the geometry of the amino acids is determined by the pre-organization of the backbone.

### Statistical analysis

An initial statistical analysis of all heme-containing protein structures from MetalPDB was conducted to establish the foundation for HemeFinder and extract the principal properties associated with heme-binding sites. The analysis focused on the three main aspects mentioned earlier: volume and shape of the binding site, the types of residues surrounding the heme-binding site, and the three geometrical descriptors (*dist(α-M), dist(β-M), angle(α-β-M)*) from the backbone that defines heme coordination. In 2022, MetalPDB reported a total of 5.164 PDB entries containing a total of 13,133 heme-binding sites. For the statistical analysis, 90% of these entries were selected randomly, while the other 10% were saved for benchmarking. However, due to poor annotations of the coordinating residues or heme name in MetalPDB or inconsistencies in PDB structures, the final number of entries analyzed was reduced to 3,812 PDB structures (9,229 heme sites) for geometrical descriptor analysis and 4,570 PDB structures (11,490 heme sites) for binding site analysis.

For the analysis of the main physical properties of the binding site, all PDB were downloaded, and each heme moiety was removed from the binding site. This analysis was performed using an in-house python script which main component is the PyKVFinder^17^, a module to define the heme cavity, obtain the volume and save all cavities as a grid of probes. From the cavity files, the shape of the binding site was characterized as an ellipsoid by its three principal axes. The environment of the binding site was defined by the different types of residues depending on their nature (**SI Figure 1**). In this analysis, all residues that were considered were situated at a distance lower than 6.5Å from carbon α and 5Å from carbon β from any point of the cavity. A second type of analysis was performed with a pychimera^18^ script to go through all the entries of the XML file from MetalPDB and calculate the three geometric descriptors and the prevalence of each coordinating residue.

This analysis revealed the average heme-binding site a volume and three principal axes values, its distribution and mean values (**Figure 2a)**. Most residues found in heme-binding sites are hydrophobic and contain large chains, being Leu and Phe the most common residues, and polar aromatic and positive residues are also quite prevalent (**Figure 2b)**. The presence of large hydrophobic residues aligns with the apolar and aromatic character of the heme, while the polar and negative nature of heme propionates can be related to the presence of polar and positive residues, especially Arg, along the absence of negative residues. The coordination sphere of the Fe and the type of heme (*a, b, c* or *d*) were also analyzed (**SI Figure 2**).

**Figure 1:**
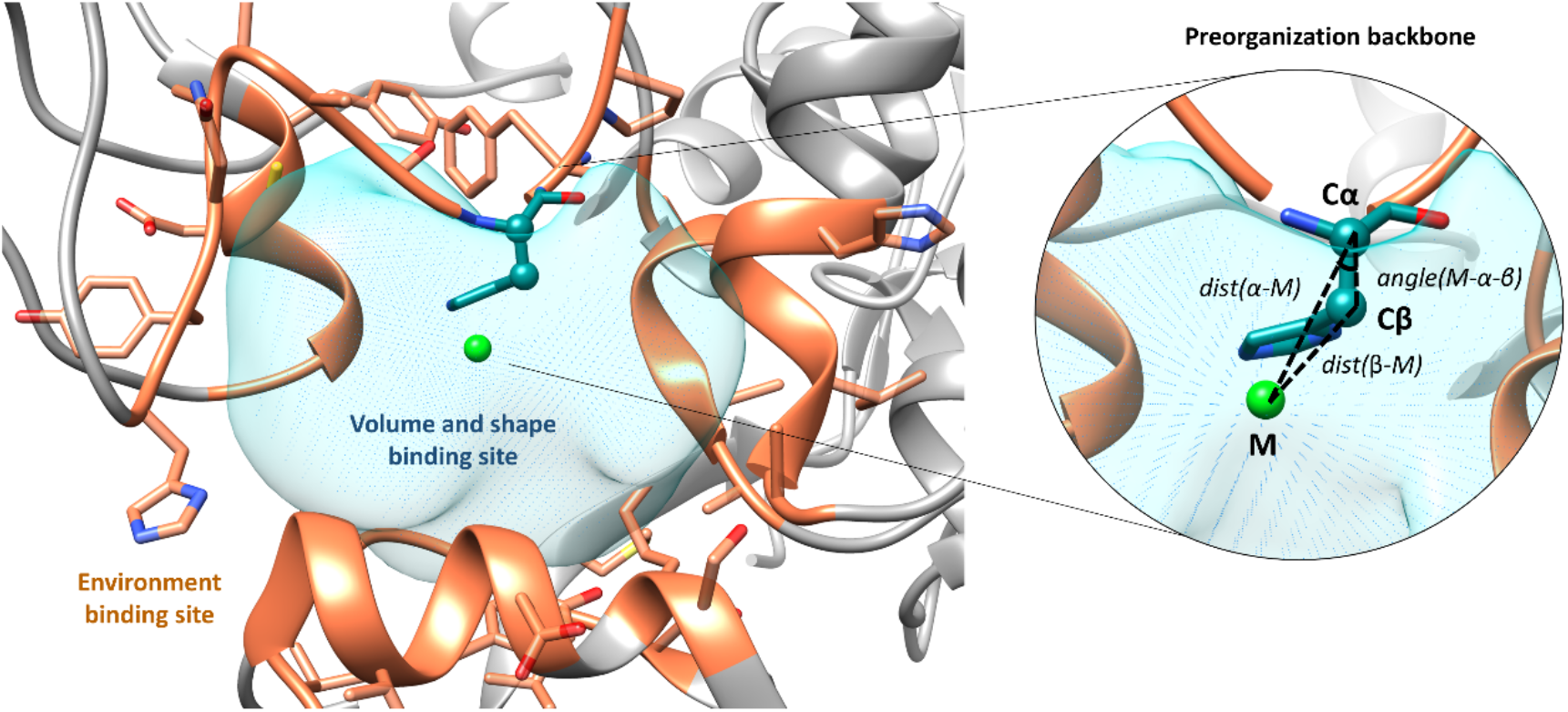
A schematic representation of HemeFinder method.

**Figure 2:**
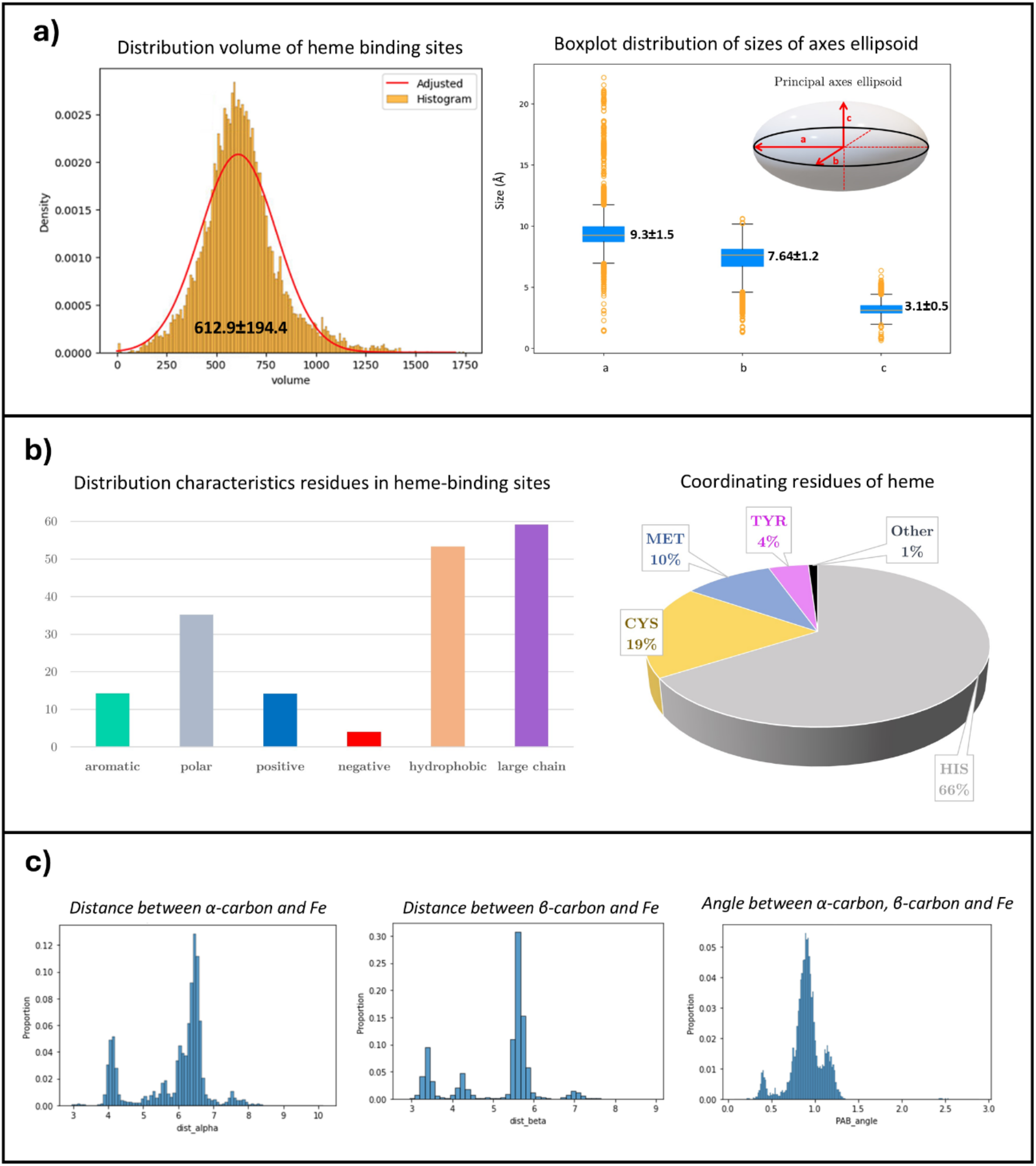
Results of statistical analysis: **a)** Distributions of volume and shape of heme-binding site **b)** Characterization of residue surrounding heme-binding sites and coordinating residues of heme. **c)** Distribution of three geometric features (dist(α-M), dist(β-M), angle(α-β-M)) that account for the preorganization of backbone for heme coordination.

The results of the second analysis show that the average coordination number to the protein is 1.35 and the most common coordinating residues are His (by far), Cys, Met and Tyr. In consequence, as default, HemeFinder considers these four main residues as coordinating and coordination number as one. This analysis only accounts for protein residues, ligands are not accounted. All three geometric features (*dist(α-M), dist(β-M), angle(α-β-M)*) are represented in **Figure 2c**. All distributions depend on the residue, but in all cases, individual residue distributions can be fitted to a bimodal function. An additional analysis was carried out for entries with two coordinating residues. Four descriptors were analyzed: the distances between α-carbons/β-carbons of both coordinating residues, the angle between α-carbons/β-carbons and the metal of both coordinating residues (**SI Figure 3 and 4**).

### Workflow

Once all the descriptors fitted from the previous steps, HemeFinder was entirely developed in Python3 with minimal dependencies. Source code, documentation, and data are available at https://github.com/laura-tiessler/hemefinder. The overall workflow of HemeFinder is divided into six different steps that are summarized graphically in **Figure 3**. The software first detects protein cavities, calculates its volume and stores them as grid points (probes). For each probe, the surrounding residues that fulfil these three geometric criteria for heme coordination dist(α-M), dist(β-M) and angle(α-β-M) are selected, with mutations suggested if possible. An ellipsoid is defined around the centroid of the probes and its size and properties are calculated to assess the suitability to bind a heme moiety. The specifics of all these steps are defined in the following section:

**Figure 3:**
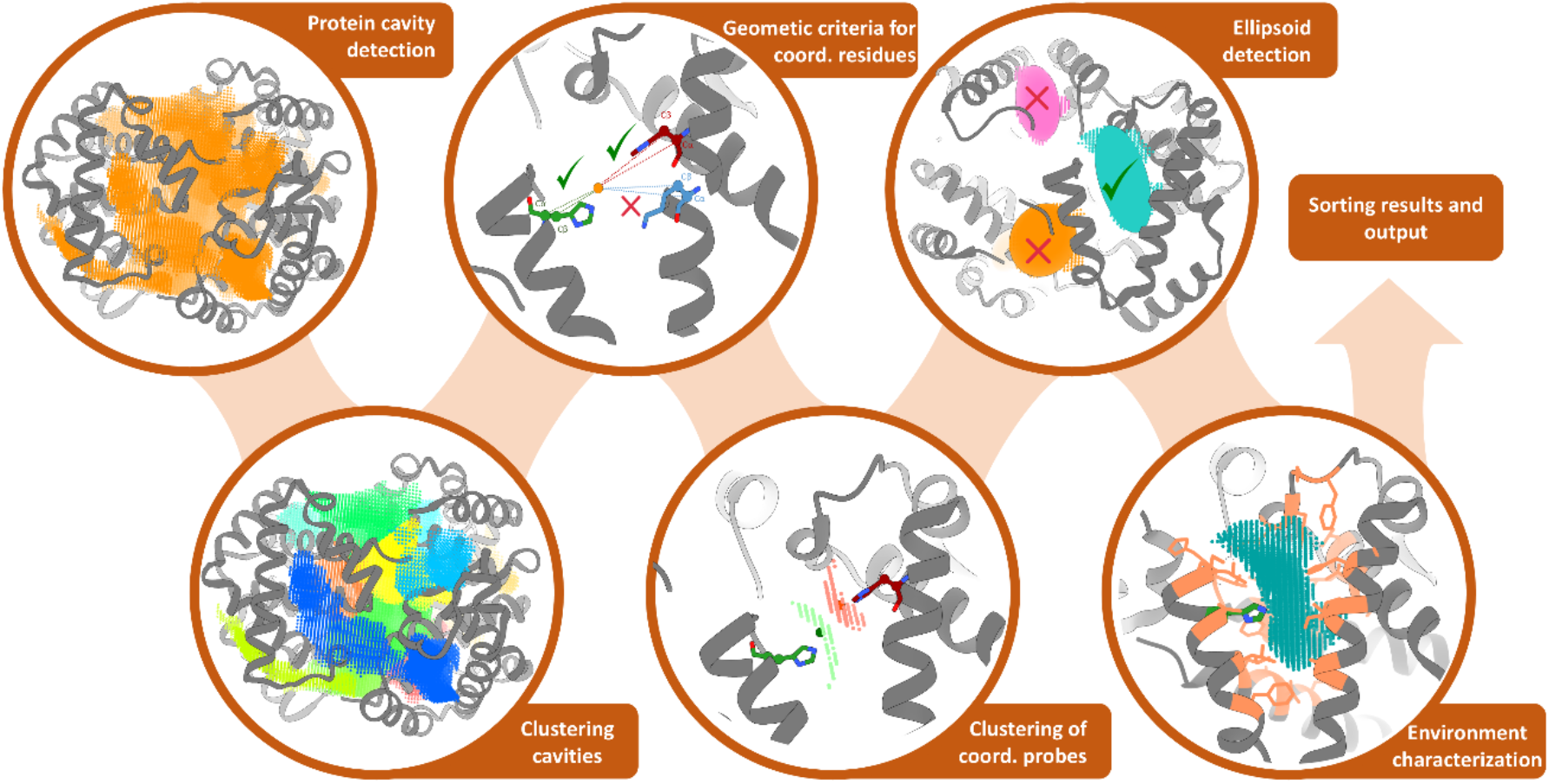
Workflow HemeFinder defined by six step process. Starting with cavity detection, followed by evaluation of geometrical criteria (dist(α-M), dist(β-M), angle(α-β-M)) to identify possible heme coordinating residues and concluding with the characterization of the heme-binding sites (ellipsoid detection and residue environment analysis).

#### Protein cavity detection

The input protein file must be provided in pdb format, or PDB ID can be specified to directly download the structure from PDB database. Protein residues are parsed to retain only coordinates and residue names of Cα and Cβ atoms. Protein cavities are detected, and their volume is calculated using the *detect* and *spatial* modules, respectively, from pyKVFinder^17^. Cavities smaller than the lower threshold established by our statistical study are discarded, as they would not be able to fit a heme molecule.

#### Clustering cavities

In large heme-binding proteins, cavities may be connected by tunnels or contain more than one heme moiety. Consequently, when the volume of the cavity is higher than the upper threshold of our statistics study, Kmeans^19^ clustering is performed to obtain the cavities of the mean size of the heme. Each cavity is saved as a grid of probes in XYZ format. Additionally, in cases where pyKVFinder leaves gaps between closely positioned residues, a function based on probe density is applied to detect these regions and fill the gaps with probes.

#### Geometric criteria

For each probe of each cavity, the software checks that *dist(α-M), dist(β-M)* and *angle(α-β-M)* are within the range that is established by our statistic study for all residues that could be coordinating. For the probes that fulfil all three criteria a scoring is assigned depending on how good the possible coordination is (explained in more details in scoring section). The output is for each probe, the coordinates of the probe, the residue number of the possible coordinating residues and its score.

#### Mutations

Based on the premise that coordination only depends on the backbone, if a residue fulfills the geometrical criteria but cannot coordinate heme, it can be mutated to one that can. If the user requests a mutation for specific residue (e.g., His or Cys), the geometric criteria is checked for all residues, not just the standard metal coordinating ones. Residues that meet the geometrical criteria for the desired mutation are considered potential coordinating residues for the next steps.

#### Clustering of probes

Once each probe has a score assigned and all the possible coordinating residues are identified, the whole set of probes are grouped in function of their coordinating amino acids, and the scores are added. This provides a list with the possible coordinating residues associated and a weighted geometric centroid with the scores.

#### Detection of the ellipsoid

From the centroid of the coordinating probes, all the probes within a sphere of size 7.63Å (determined from statistic study) are selected. These probes are modelled into an ellipsoid, characterized by its three axes and center. If the three axes fall below the threshold of the statistical study, the result is discarded, as this indicates that the heme would not be able to bind fit in the hypothetical binding site. For each ellipsoid, a score is assigned depending on the alignment of the centroid and the ellipsoid center.

#### Residue environment

a distance matrix is calculated between all the probes of the ellipsoid and the α-carbon and β-carbon from all residues that predicted cavity. Residues with α-carbon and β-carbon within 6.5 Å and 5.5 Å from an ellipsoid probe, respectively, are stored. The proportion of each type of residue is calculated and use as a score that represents how adequate the environment of the ellipsoid is (further explained in the scoring section).

#### Sorting and scoring

Results from different clusters are combined. For results where there are two possible coordinating residues, an additional geometrical check is performed. The distances and angles between the Cα, Cβ and the centroid probe must be within the statistical thresholds. If not, it is discarded as the putative coordinating residues are out of the range for coordination. Finally, coordination, ellipsoid and residues scores are normalized and the final score the sum of these normalized scores, ranging between 0 and 3, with 3 being the highest score possible.

#### Output

The output of the HemeFinder is a pdb file (using different atom types) that includes the centroid of the probes (He), all the probes that make up the ellipsoid (Xe) and the coordinating probes (Ne) (**Figure 4**). Additionally, the software prints all the results and exports them into a json file, which contains all the possible heme-binding sites sorted by score and with its corresponding coordinating residues.

**Figure 4:**
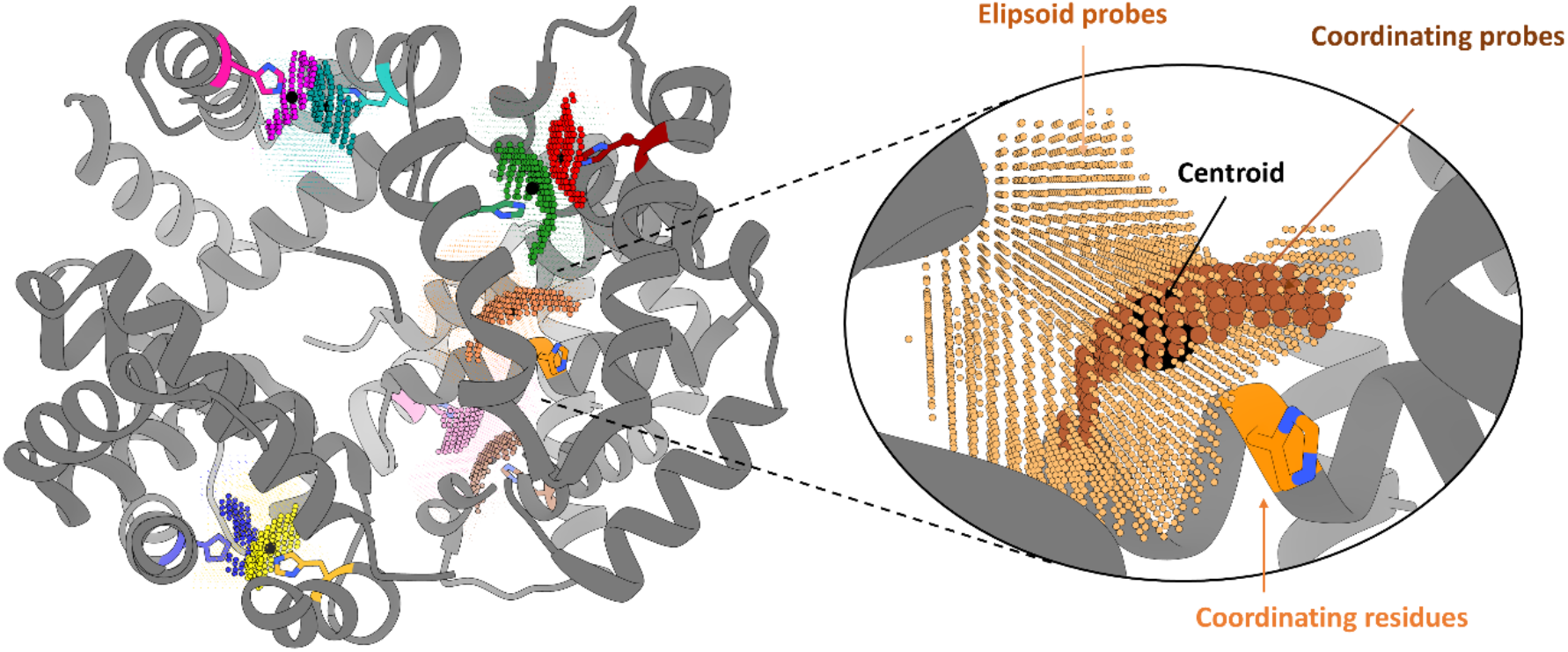
Example of output from HemeFinder. Each result is color-coded, displaying the ellipsoid of the possible heme characterized by probes, the coordinating probes and centroid representing the position of Fe.

### Scoring

The scoring of HemeFinder aims to assess the suitability of a cavity for the binding of the heme considering three different items: geometrical criteria of backbone, residue composition of environment and metal centrality in ellipsoid. Here we focus on the main aspects of the scoring we established but further information can be found in ESI.

#### Geometrical criteria of backbone

The analysis of three geometrical descriptors *dist(α-M), dist(β-M)*, and *angle(α-β-M)-* was performed to obtain fitness function that measures how close is the preorganization of certain residues to the ideal geometry for binding of heme. The distribution of each geometrical descriptor was approximated as a bimodal probability density function, chosen for its simplicity and for fitting the observed data (**SI Figure 5**). Therefore, the likelihood that a given residue will be able to coordinate given a set of geometrical parameters *x* is approximated as the union of the three distributions as follows:

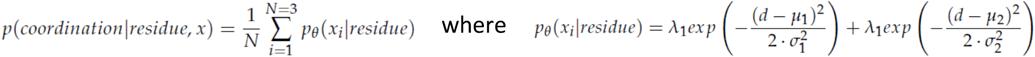

Furthermore, it is expected that some coordinating residues have more tendency to coordinate heme than others. This is modeled as the proportion of coordinations that each residue establishes in the database over the total number of coordinations documented for heme. This allows coordinations involving a more common coordinating residue to be scored higher than coordinations involving fewer common residues.

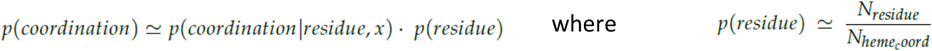

#### Residue composition

In addition to the preorganization of the backbone, the residue composition around the heme is critical for its binding. This scoring function assesses the residue environment around the potential heme-binding site. The statistical analysis determined the most optimal environment for stabilizing the heme group by dividing the residues into 5 groups depending to their chemical nature (**SI Figure 1**) and calculating its proportions. The density function for each of the distributions generated was approximated with a bimodal function as in the geometric criteria (**SI Figure 5**). Thus, given a set *c* of features of the chemical environment of a possible heme-binding site, the probability of that environment being suitable for heme-binding is approximated as a sum of the probabilities for each of the five types of residues:

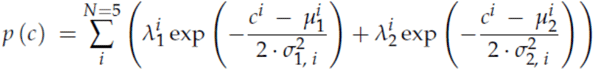

#### Metal centrality in ellipsoid

The aim of this score is to measure the centrality of the calculated centroid of the coordinating probes against the ellipsoid, which will give an indication of how deviated from the center of the ellipsoid is the centroid. This is modelled as the distance between the centroid of the coordinating probes (C_p_) and the center of the ellipsoid (C_e_) divided by the mean length of the three axes of the ellipsoid (a, b, c).

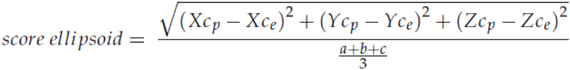

All residue distributions fitted to the bimodal functions for the geometrical criteria and residue composition found in **SI Figure 6 and 7**.

## RESULTS AND DISCUSSION

To evaluate the quality of HemeFinder to predict heme-binding sites, we carried out two benchmarks. Furthermore, to better illustrate the applicability of the software we also provide with a case study where HemeFinder is bridged with protein-ligand dockings.

### Benchmarks

Benchmark 1 contains 474 proteins, which correspond to the remaining 10% of PDB entries from the MetalPDB that were not included in the training set of HemeFinder. Benchmark 2 includes all the structures of heme containing systems added to the Protein Data Bank until the date of writing of the manuscript^3^. The set was filtered to avoid protein redundancy, resulting in a total set of 56 unique proteins. Both datasets were prepared by downloading all structures from the PDB, removing all cofactors and solvent and selecting only one monomeric structure. HemeFinder was applied to all systems with default parameters (**SI Figure 8**).

Results of benchmark 1 and 2 present success rates of approximately 94% when considering that the any of the solutions present the correct identification of the crystallographic heme-binding sites (**Table 1**). When considering also that the the exact coordinating residues of the iron (one or multiple coordinating residues), the success rate only decreases slightly (approximately 91, **Table 1**). Similarly to docking approaches, the performance of HemeFinder can also be evaluated through its ranking accuracy, the number of solutions of which % was found in the top 1, top 3, or top 10 predictions. For entries containing more than one heme-binding site, the average rank was considered. The average rank position is between 2 and 3 in both benchmarks and in general between 10 and 20 solutions are obtained for each run. Though finding the best solution (ranked top 1) reduces the apparent predictiveness (ca. 40% in average), analyzing 3 or 10 solutions already shows very accurate prediction of heme-binding sites (above 80 and 90% respectively). Those results are more than promising for any prediction. Moreover, the average computation time was 52s per entry, or 12s if large proteins are excluded. This speed makes HemeFinder an excellent tool for high-throughput screenings. Results for Benchmark 2 are shown as an example in **SI Figure 9**.

**Table 1:** Results obtained for Benchmark 1 and 2. Success rate, ranking of solutions (top 1, top 3 and top 10), average rank of solution and average number of solutions are represented.

|  | Success rate | Ranking top 1 | Ranking top 3 | Ranking top 10 | Average rank solution | Average num. solutions |
| --- | --- | --- | --- | --- | --- | --- |
| **Benchmark 1** | 94.3<br>(91.77) | 48.73<br>(38.82) | 81.01<br>(72.57) | 94.94<br>(90.72) | 2.51<br>(5.47) | 11.4 |
| **Benchmark 2** | 94.64<br>(91.07) | 41.07<br>(32.14) | 82.14<br>(66.07) | 94.64<br>(89.29) | 2.83<br>(6.22) | 21.57 |

The success rate that HemeFinder shows corresponds to current state-of-the-art programs. Moreover, the philosophy of HemeFinder is substantially different from the currently available heme binding site predictor. Still, it is important to highlight that the goal of HemeFinder is not to find heme-binding sites only, but to find protein cavities sites that have the convenient geometrical and physicochemical requisites to become heme-binding compatible. Because of this, one of its byproduct, is to allow the identification of suboptimal binding sites which could be associated to transient sites that could be involved in possible heme pathways. To better demonstrate those concepts and the strength of HemeFinder in this context, we decide to deepen into one particular illustrative case.

### Detection of heme-binding sites in Heme Carrier Protein 1

To show the interest of HemeFinder in detecting heme-binding sites in protein structures, we focused on the Proton-Coupled Folate Transporter (PCFT) also identified as the Heme Carrier Protein 1 (HCP1). This transporter was originally identified as a heme carrier and found in the small intestine, specifically in the duodenum.^20^ However, it is was latter characterized that it acts as a proton-coupled folate transporter.^21,22^ The debate between both functions was finally resolved by identifying a dual function with higher affinity for folate and low affinity for the heme (K_M_=1.67 µM and 125 µM respectively).^23^ Therefore, the current hypothesis is that the receptor acts primarily as a folate transporter, but also transport the heme more reversibly. Although, heme interaction with PCFT/HCP1 has clearly been demonstrated experimentally, one critically missing piece of information is the lack of heme-bound structure. This is fundamental for understanding the full mechanism of heme uptake. Therefore, where and how the heme could bind to the receptor is unknown.^24,25^

So far, only one structure has been determined for HCP1, which was obtained by cryo-EM from *Gallus gallus* (PDB 7BC7).^26^ This structure is in its apo form and in open conformation towards the extracellular site of the transporter (HCP1-extra). Additionally, an AlphaFold structure is available with open conformation towards the cytosolic site (HCP1-cyto). Comparison of both structures is depicted in **SI Figure 10**. The objective of this part of the work is to use HemeFinder to determinate where the heme group can bind, which would be the path of heme entrance in both protein conformations and with which residues would coordinate the heme. HemeFinder was applied using default parameters on both the X-ray (open) and theoretical (closed) structures (HCP1-extra and HCP1-cyto).

HemeFinder calculations reveal that, in the open conformation HCP1-extra, one possible binding site appears in between His70 and Tyr422, whereas in the open conformation HCP1-cyto there are two possible binding sites with interaction with His255 and Cys405 respectively (**Figure 5a**). These two heme-binding regions would correspond to entrance or exit path for each site of the transporter. However, calculations in both conformations suggest the same heme-binding regions on the center of the transporter. There are three main binding heme regions in the center of the helix barrel: **1)** region of Tyr165 and His289, best scored in both systems, **2)** region in which heme could coordinate with either Tyr296 or Tyr320, **3)** in HCP1-cyto, there is a site that has good score in conformation open towards cytosol, in which heme could coordinate with Tyr323. Apart from providing the first insights on the heme-binding sites of HCP1, one of the more relevant results of the HemeFinder calculations is to reveal that most heme coordinating residues would correspond to Tyr, which are classified as a relatively low energy binder to the iron when compared to other residues like histidine or cysteine. This could explain the low affinity for heme of this receptor and more particularly. The residues involved in the heme pathway and its scores in each form are shown in **Figure 5b**.

**Figure 5:**
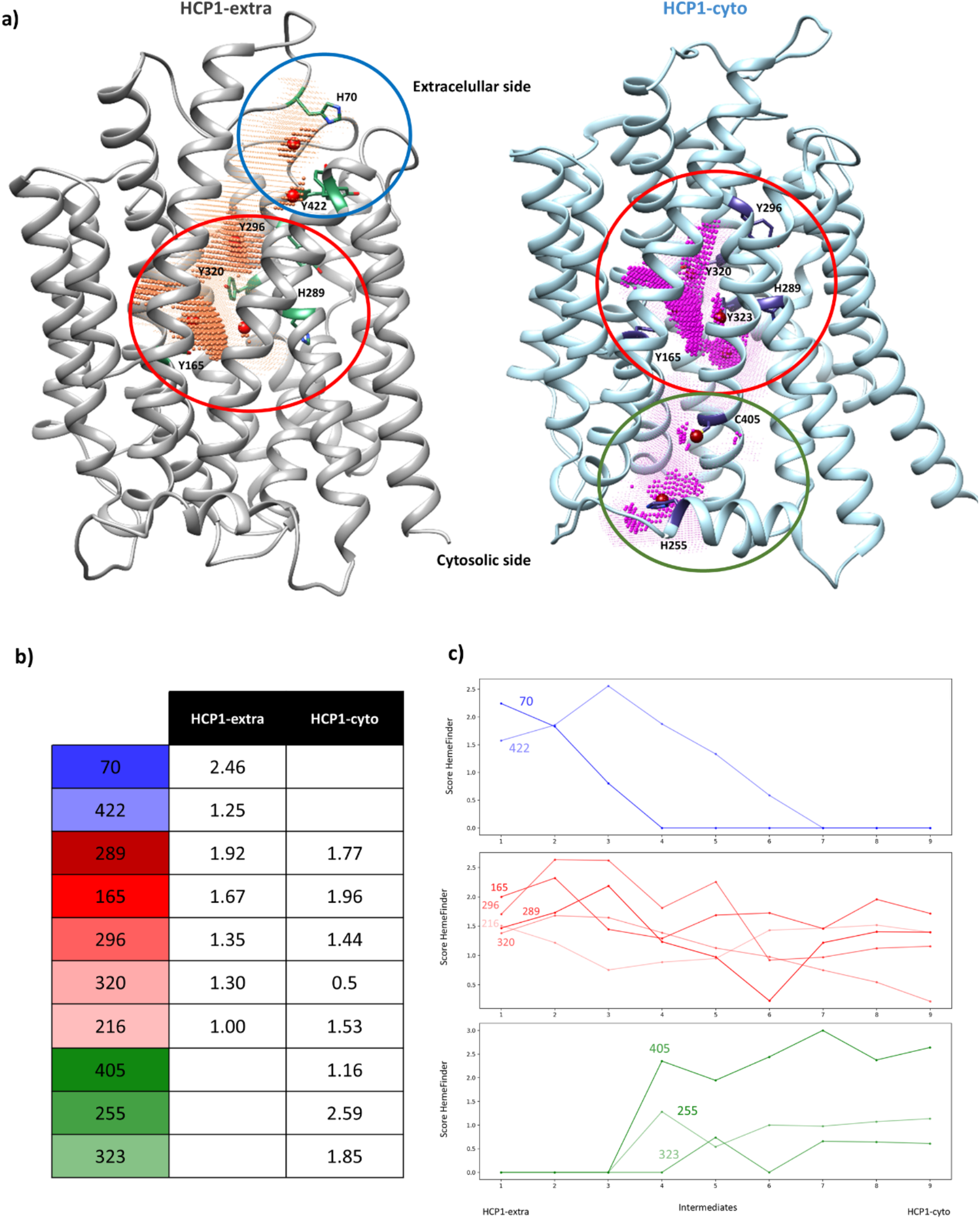
**a)** HemeFinder results obtained for HCP1 in open conformation toward extracellular space (HCP1-extra in gray) and HCP1 in open conformation toward cytosol (HCP1-cyto in blue) with possible heme coordinating residues highlighted. **b)** Scores from all possible heme coordinating residues obtained by HemeFinder. **c)** Representation of score each intermediate over intermediates HCP1-extra (1) and HCP1-cyto (9).

To improve the conformational sampling of the protein for these exercises, we used the Morphing tool implemented in UCSF Chimera.^27,28^ Nine intermediate states between both conformations were obtained and HemeFinder was used to find heme-binding sites for all intermediates. (**Figure5c)**. The scoring for each binding site is depicted along the intermediates, number one representing the closest to HCP1-extra and number 9 to HCP1-cyto. In this representation it can be observed how His70 and Tyr422 disappear as the structures move towards the open cytosolic form (blue), whereas His225, Tyr323 and Tyr405 start to appear (green). The binding sites found in the center of the transporter (in red) are maintained in all states, but their scoring fluctuates in the different intermediate states.

This clearly shows that this central region corresponds to the heme-binding site and there are several possible coordinating residues, with two potential pathways, 70-422 and 255-405, that lead to it. To validate these solutions, molecular dockings were performed to confirm that heme is able to bind in all these regions. Dockings reveal that in some cases Tyr coordination could not be observed, maybe due to rotamer restrictions. However, they also show that both Tyr422 in the extracellular site and His255 in the cytosolic site would be able to bind heme and coordinate it as the major transient sites between the core of the protein and the intra- or extracellular media. The accuracy of HemeFinder combined with its velocity allows us to identify both the main heme-binding site of the transporter and possible routes for the crossing of the member from extra to intra-cellular media.

## CONCLUSIONS

Despite the importance of heme, the binding of heme to proteins and its prediction has not been widely studied. So far, most programs are based on sequence-based predictions which could be valuable for proteins from the natural realm but could be limitative for de novo ones. In this work we present HemeFinder, a new program that allows detecting heme-binding sites and designing new hemo-enzymes based only on the protein structure, the composition of the possible binding cavities and the geometrical predisposition of the backbone.

HemeFinder has been able to predict more than 94% of crystallographic structures from benchmark, ranking possible heme-binding sites considering how preorganized the backbone of the coordinating residues, the characteristics of the environment and the shape of the binding site. HemeFinder shows great potential for detecting pathways of natural heme-binding sites, but also for the design of new ArMs. For example, due to the high speed of the program it can be used to screen possible scaffolds for the design of ArMs. Compared to other software for heme prediction, HemeFinder offers a fast approach to detect heme-binding sites that can be further validated by molecular dockings, or it could even be combined with other docking tools.

Combined with X-ray structure, AlphaFold model and docking approaches, Hemefinder has also shown to be a promising tool in elucidating heme-binding mechanism on system orphan of molecular knowledge. The application on HCP1, the unique heme transporter characterized in the duodenum in most evolved organisms, the speed and versatility of Hemefinder allows to identify the main binding site at the core of the protein as well as possible transient sites at the extreme of the membrane regions.

Altogether, Hemefinder appears a novel tool with high potentiality in the fields of research involving heme-binding mechanism.

## Acknowledgments

All authors acknowledge the support of the Generalitat de Catalunya (2021 FISDU 00019) and the Spanish Ministerio de Ciencia e Innovación for Grant PID2020 116861GB-I00 and PID2023 149492NB-I00. L. T.-S. thanks the Spanish MINECO for grant FPU18/05895.

## Footnotes

1 December 2023

