## Supporting information for "HemeFinder: a Computational Predictor for Heme-Binding Sites in Proteins"

**Aromatic**: PHE, TYR, TRP
**Polar**: TYR, THR, SER, CYS, MET, ASN, GLN, HIS,
**Positive:** HIS, LYS, ARG,
**Negative:** ASP, GLU,
**Hydrophobic:** PHE, TRP, ALA, VAL, LEU, ILE, PRO,
**Large chain**: PHE, TYR, TRP, MET, GLN, GLU, ARG, LYS, HIS, LEU, ILE

**SI Figure 1:** Classification residues of residues depending on their chemical nature for residue composition of heme binding environment.

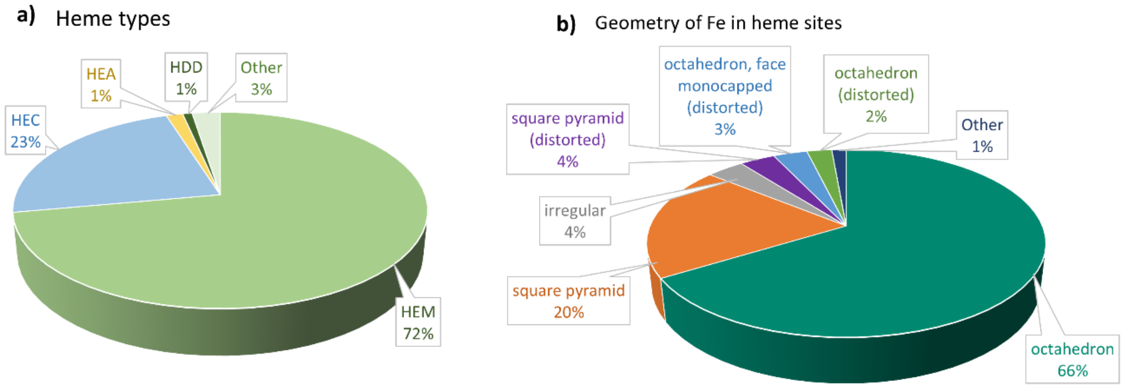


**SI Figure 2: a)** Results statistical analysis of most common heme types (HEA=heme a, HEM=heme b, HEC = heme c and HDD=heme d) in dataset. **b)** Results analysis geometry of heme binding sites in dataset.

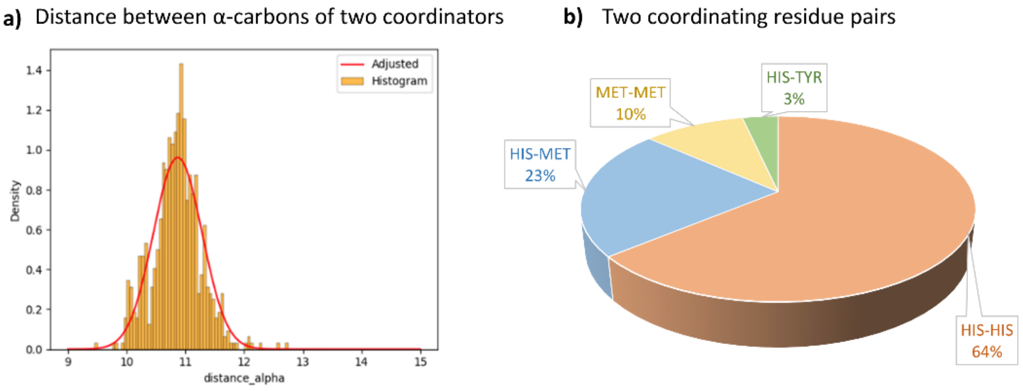


**SI Figure 3: a)** Distribution of distances between C-$\alpha$ of two coordinating residues **b)** Proportion of two coordinating residues.

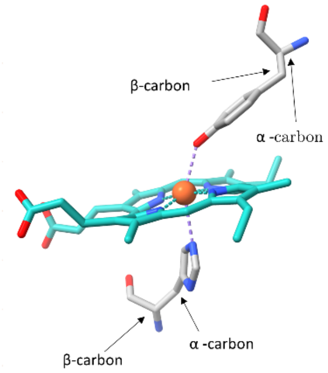


| | Angle metal α-carbons | | Angle metal β-carbons | | Distance α-carbons | | Distance β-carbons | |
|---|---|---|---|---|---|---|---|---|
| | min | max | min | max | min | max | min | max |
| ['CYS', 'HIS'] | 138,01 | 158,56 | 132,81 | 196,59 | 9,45 | 11,43 | 8,41 | 9,69 |
| ['HIS', 'HIS'] | 121,90 | 179,91 | 134,60 | 179,92 | 11,00 | 13,57 | 10,33 | 11,63 |
| ['HIS', 'LYS'] | 101,08 | 190,47 | 114,71 | 188,82 | 9,82 | 16,64 | 9,16 | 13,80 |
| ['HIS', 'MET'] | 97,16 | 173,84 | 125,03 | 172,26 | 9,83 | 11,91 | 8,45 | 10,37 |
| ['HIS', 'TYR'] | 104,05 | 177,12 | 82,89 | 181,22 | 8,98 | 17,53 | 9,05 | 12,84 |
| ['MET', 'MET'] | 156,67 | 168,99 | 170,58 | 183,67 | 10,67 | 11,57 | 8,07 | 8,99 |
| ['MET', 'TYR'] | 164,78 | 175,14 | 154,55 | 173,27 | 13,26 | 14,27 | 10,79 | 11,76 |

**SI Figure 4:** Results analysis for entries dataset with two coordinating residues. Results include the distances between α-carbons/β-carbons of both coordinating residues, the angle between α-carbons/β-carbons and the metal of both coordinating residues

$$p_\theta(x_i|residue) = \lambda_1 exp\left(-\frac{(d-\mu_1)^2}{2\cdot\sigma_1^2}\right) + \lambda_1 exp\left(-\frac{(d-\mu_2)^2}{2\cdot\sigma_2^2}\right)$$

$$p\,(c) \;=\; \sum_{i}^{N=5}\left(\lambda_1^i \exp\left(-\frac{c^i \;-\; \mu_1^i}{2\cdot\sigma_{1,\,i}^2}\right) + \lambda_2^i \exp\left(-\frac{c^i \;-\; \mu_2^i}{2\cdot\sigma_{2,\,i}^2}\right)\right)$$

**SI Figure 5:** The score of the residue is defined by a bimodal distribution with a set of 6 parameters $\theta = \lambda_1, \lambda_2, \mu_1, \mu_2, \sigma_1, \sigma_2$. $\mu_i$: are the centers of the distributions; $\sigma_i$: are the deviations; $\lambda_i$ a correction factor that accounts for the relative contributions of each component of the distribution. To find the most appropriate set of parameters θ, the empirical data distributions were smoothen with a Kernel Density Estimation (KDE) with gaussian kernel and then the curves $p_\theta$ $(x_i|\text{residue})$ were adjusted to the smooth distribution using the Levenberg-Marquadt algorithm for non-linear least-squares adjustment.

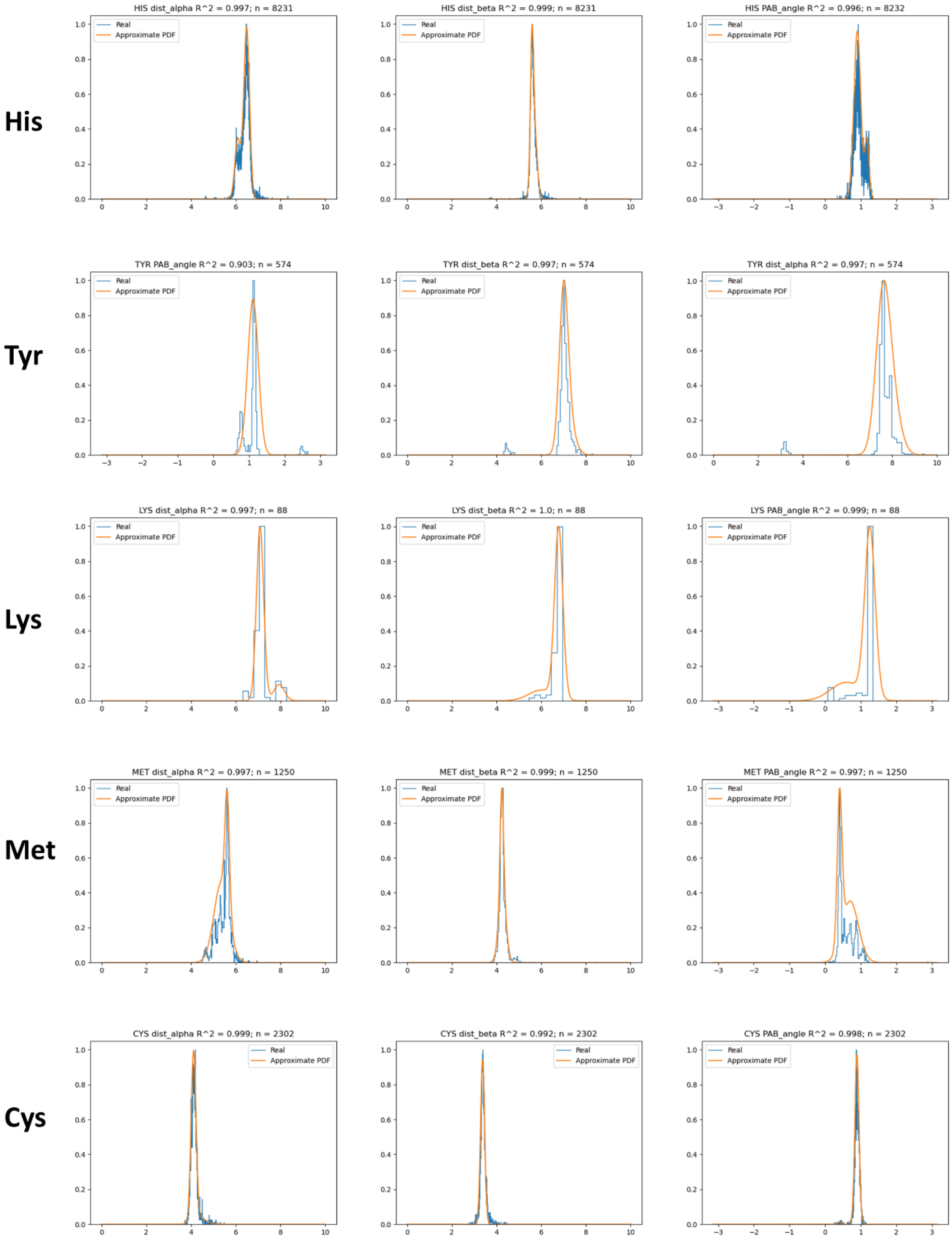


**Figure 6:** Bimodal distribution fitting for His, Tyr, Lys, Met and Cys for three geometrical criteria (dist(*α*-M), dist(β-M) and angle(*α*-β-M)).

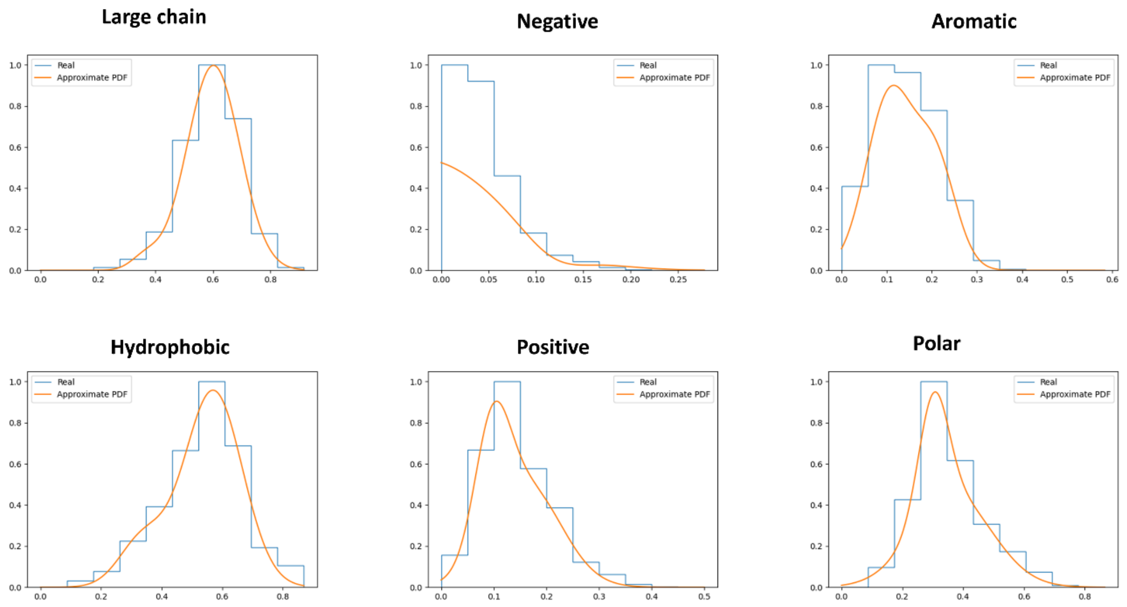


**Figure 7:** Bimodal distribution fitting for types of residues found in heme-binding sites.

--**output:** Directory where outputs should be stored.
--**coordinators:** List of possible coordinating residues
--**mutations:** List of possible mutating residues
--**num_coordinants:** List of possible mutating residues
--**probe_in:** Probe in size for pyKVFinder cavity calculation
--**probe_out**: Probe out size for pyKVFinder cavity calculation
--**removal_distance:** removal distance for pyKVFinder cavity calculation
--**volume_cutoff:** Volume cutoff for pyKVFinder cavity calculation
--**surface**: SES or SASA for pyKVFinder cavity calculation

**Default parameters**

**coordinators**=HIS, TYR, CYS, MET
**Mutations =** 0
**num_coordinants** = 1
**probe_in** = 1.5
**probe_out** = 11.0
**removal_distance** = 2.5
**volume_cutoff** = 1.5
**surface** = "SES"

**Figure 8:** Possible inputs and default parameters for HemeFinder.

| PBD | Binding site rank | Number solutions |
|---|---|---|
| 6iss | 2 | 4 |
| 6z1m | 1 | 14 |
| 6kai | 1 | 2 |
| 7cjj | 40 | 139 |
| 6kmn | 1 | 6 |
| 6ot7 | 4 | 7 |
| 6kmk | 2 | 7 |
| 7mhy | 0 | 16 |
| 7y44 | 3 | 22 |
| 7zxy | 8 | 37 |
| 7cub | 5 | 39 |
| 6y2y | 1 | 8 |
| 6bww | 1 | 3 |
| 6myo | 9 | 24 |
| 6ay4 | 7 | 17 |
| 6m8f | 1 | 3 |
| 7n3l | 2 | 5 |
| 6i7c | 1 | 3 |
| 6te7 | 1 | 4 |

| PBD | Binding site rank | Number solutions |
|---|---|---|
| 6pqw | 2 | 2 |
| 7kqr | 1 | 11 |
| 7s3z | 2 | 12 |
| 7dvt | 4 | 6 |
| 7ou5 | 1 | 6 |
| 7vn0 | 2 | 15 |
| 6gd6 | 5 | 9 |
| 7rh5 | 21 | 117 |
| 6c3h | 2 | 2 |
| 6i7z | 1 | 6 |
| 5zkv | 2 | 7 |
| 7vuc | 1 | 4 |
| 6i3t | 2 | 4 |
| 7s9y | 5 | 40 |
| 7lq5 | 11 | 34 |
| 7su2 | 0 | 6 |
| 6k3q | 2 | 5 |
| 6fl2 | 1 | 3 |
| 7acp | 1 | 4 |

| PBD | Binding site rank | Number solutions |
|---|---|---|
| 6k3o | 6 | 7 |
| 7bok | 1 | 8 |
| 7ant | 2 | 5 |
| 7s3t | 1 | 2 |
| 6j7i | 1 | 33 |
| 7tsa | 3 | 9 |
| 6t15 | 10 | 364 |
| 6ng1 | 4 | 9 |
| 6rtd | 2 | 19 |
| 6rte | 2 | 18 |
| 6t6v | 4 | 26 |
| 7s3x | 2 | 14 |
| 6cre | 2 | 2 |
| 6pyz | 1 | 11 |
| 6vjx | 2 | 2 |
| 6r1q | 3 | 5 |
| 6pn0 | 2 | 11 |
| 6m4q | 3 | 10 |

**SI Figure 9:** Results obtained in benchmark 2 including the ranking for the first solution in the heme-binding site of the crystallographic structure and the number of total solutions.

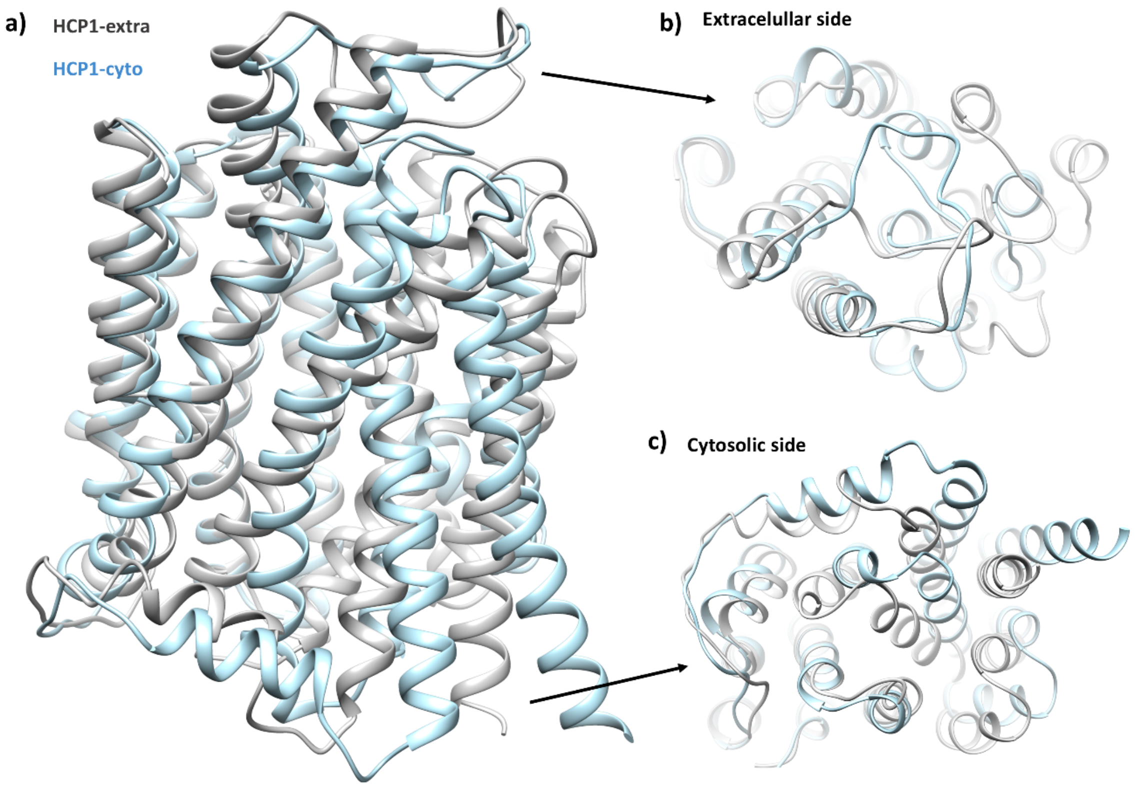


**SI Figure 10:** Superposition of structures HCP1-extra (gray) and HCP1-cyto (blue) from **a)** a lateral view **b)** the extracellular side and **c)** the cytosolic side.
